## Supplemental Figures for "Releasing a preprint is associated with more attention and citations for the peer-reviewed article"

Figure S1

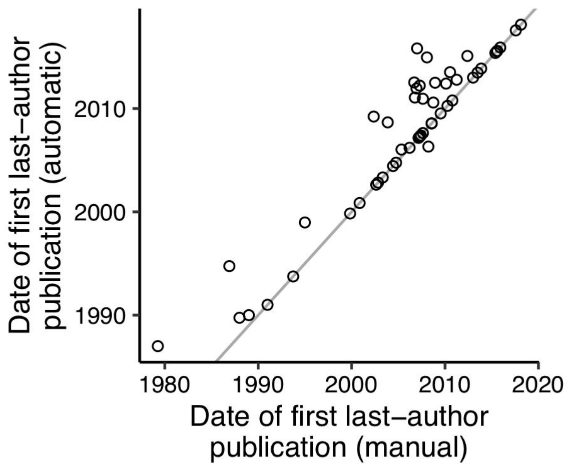


Accuracy of automatically inferring last-author publications from names and affiliations in PubMed. Each point represents one of the 100 randomly selected articles. The gray line represents y = x. For details, see Table S2.

### Figure S2

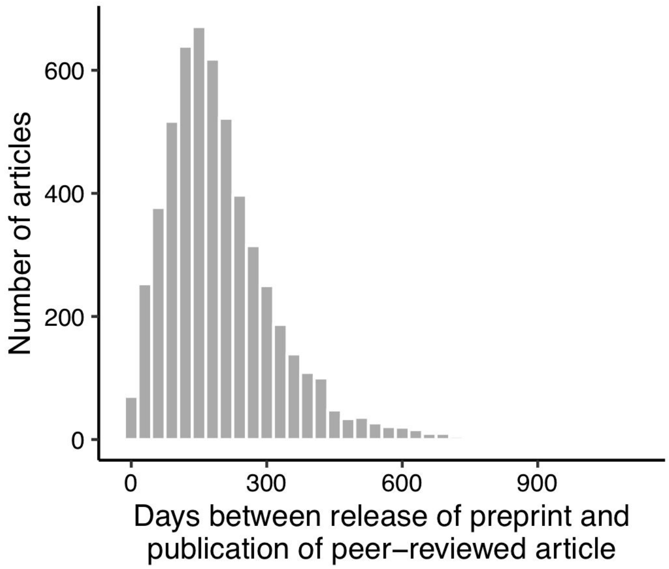


Histogram of the number of days by which release of the preprint preceded publication of the peer-reviewed article, including articles from all journals.

Figure S3

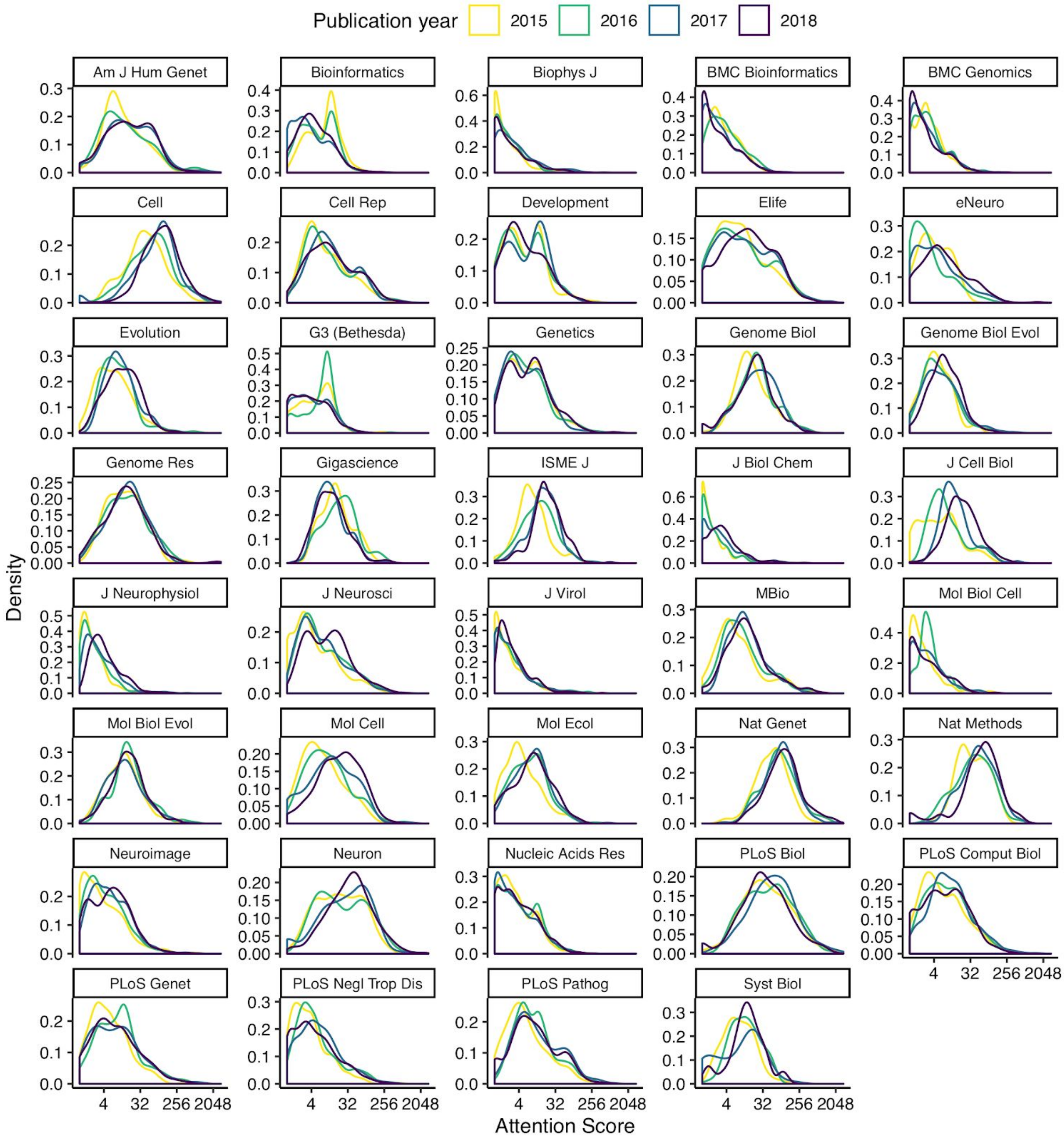


Kernel density estimates of Attention Score (with a pseudocount of 1) for articles in each journal and each publication year. Estimates were computed using the default settings of the geom_density function of the ggplot2 R package.

Figure S4

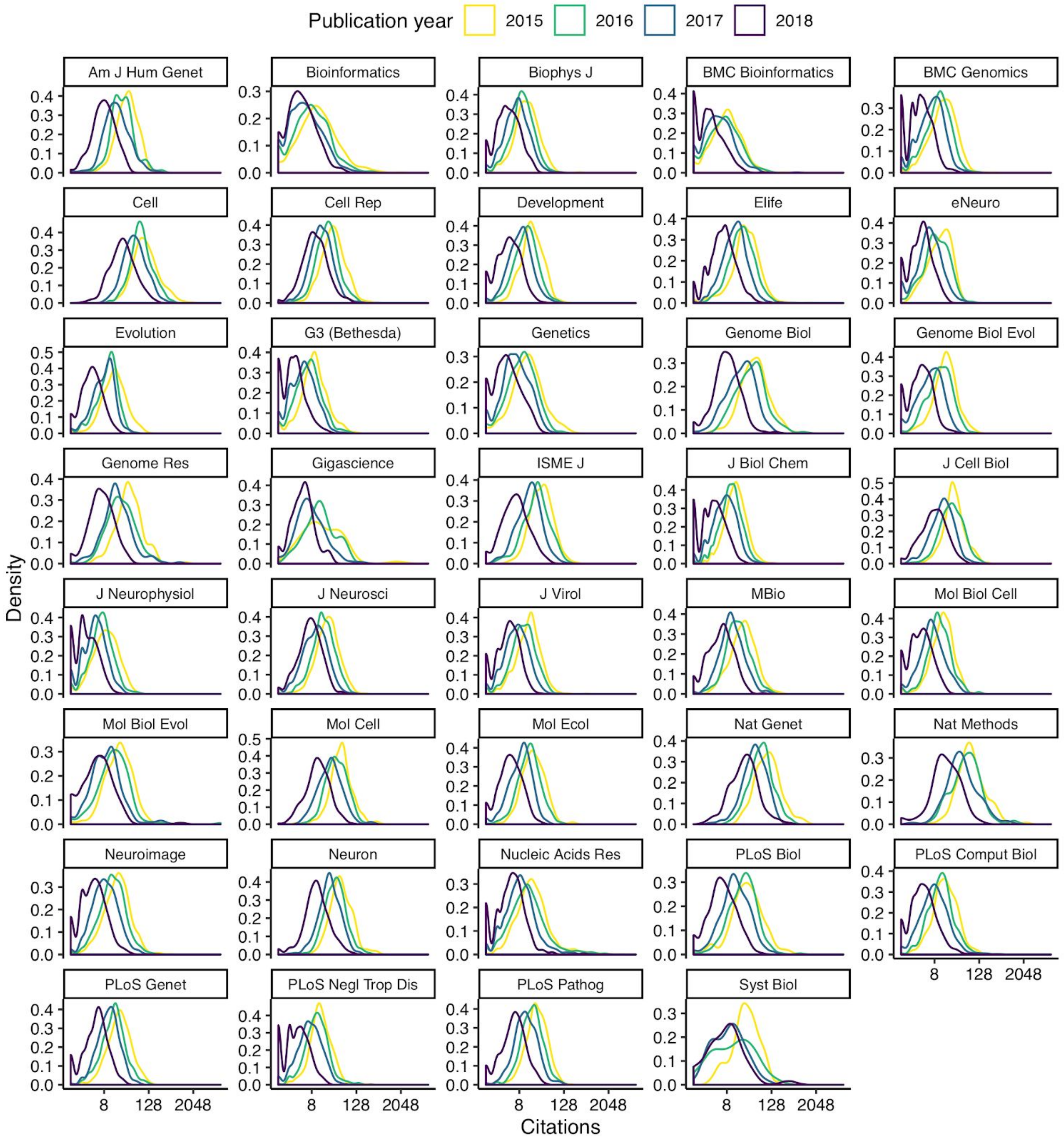


Kernel density estimates of number of citations (with a pseudocount of 1) for articles in each journal and each publication year. Estimates were computed using the default settings of the geom_density function of the ggplot2 R package.

Figure S5

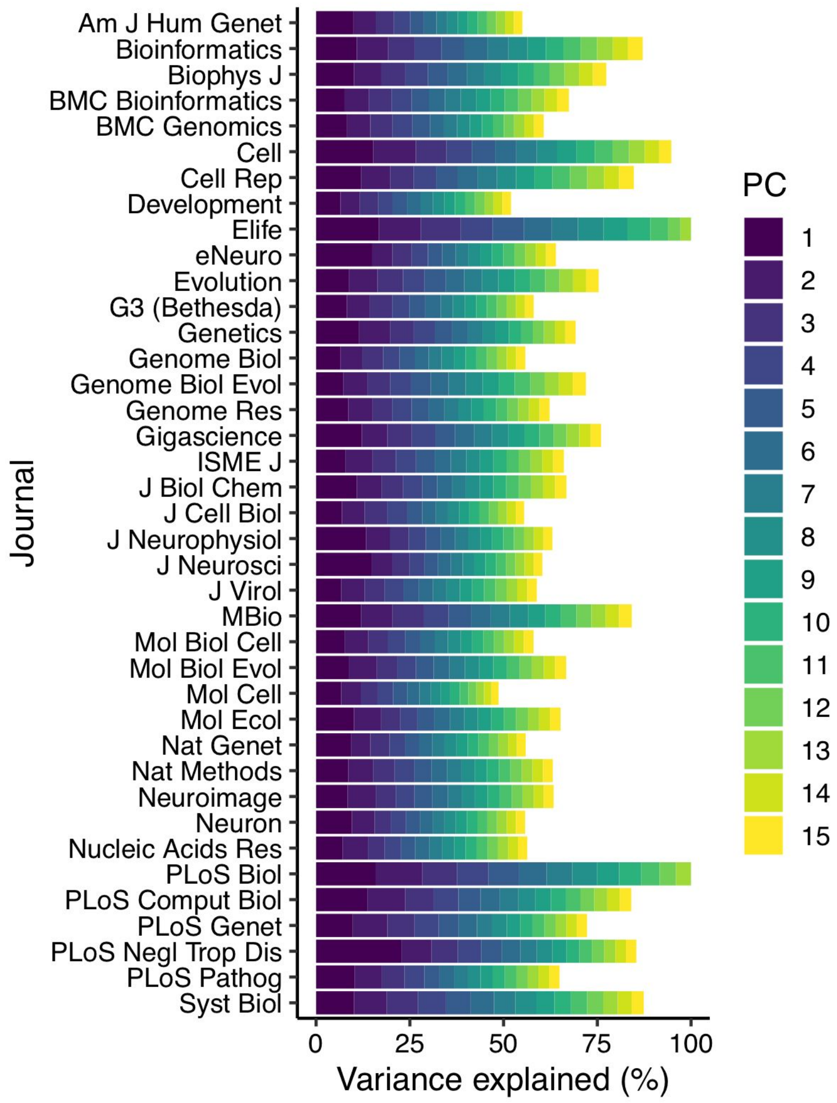


Percentage of variance in MeSH term assignment explained by the top 15 principal components for each journal.

Figure S6

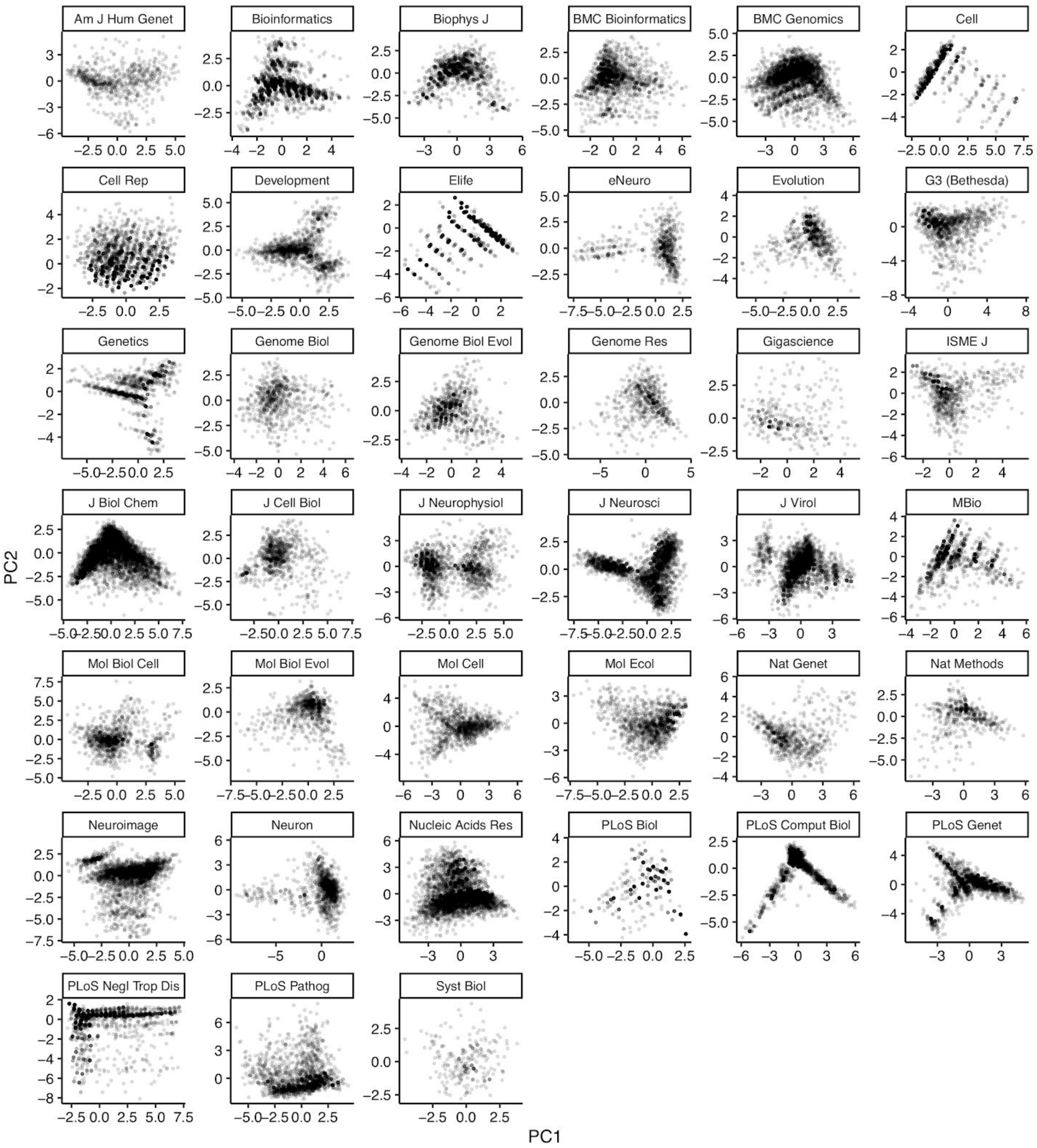


Scores for the top two principal components of MeSH term assignments for each journal. Each point represents an article.

Figure S7

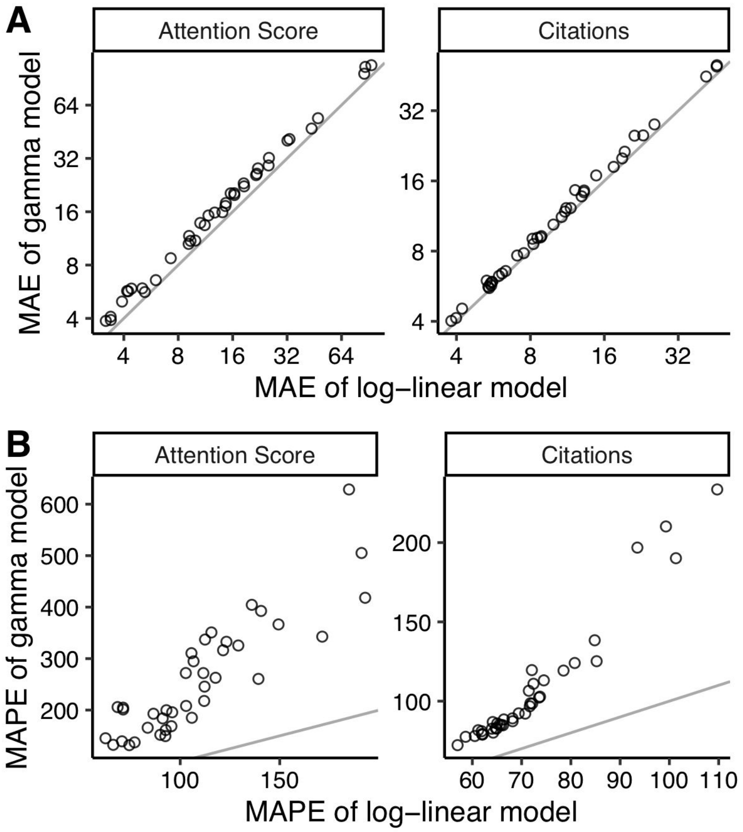


Comparing mean absolute error (MAE) and mean absolute percentage error (MAPE) of Gamma and log-linear regression models for each metric. Each point represents a journal. The gray line indicates y = x.

Figure S8

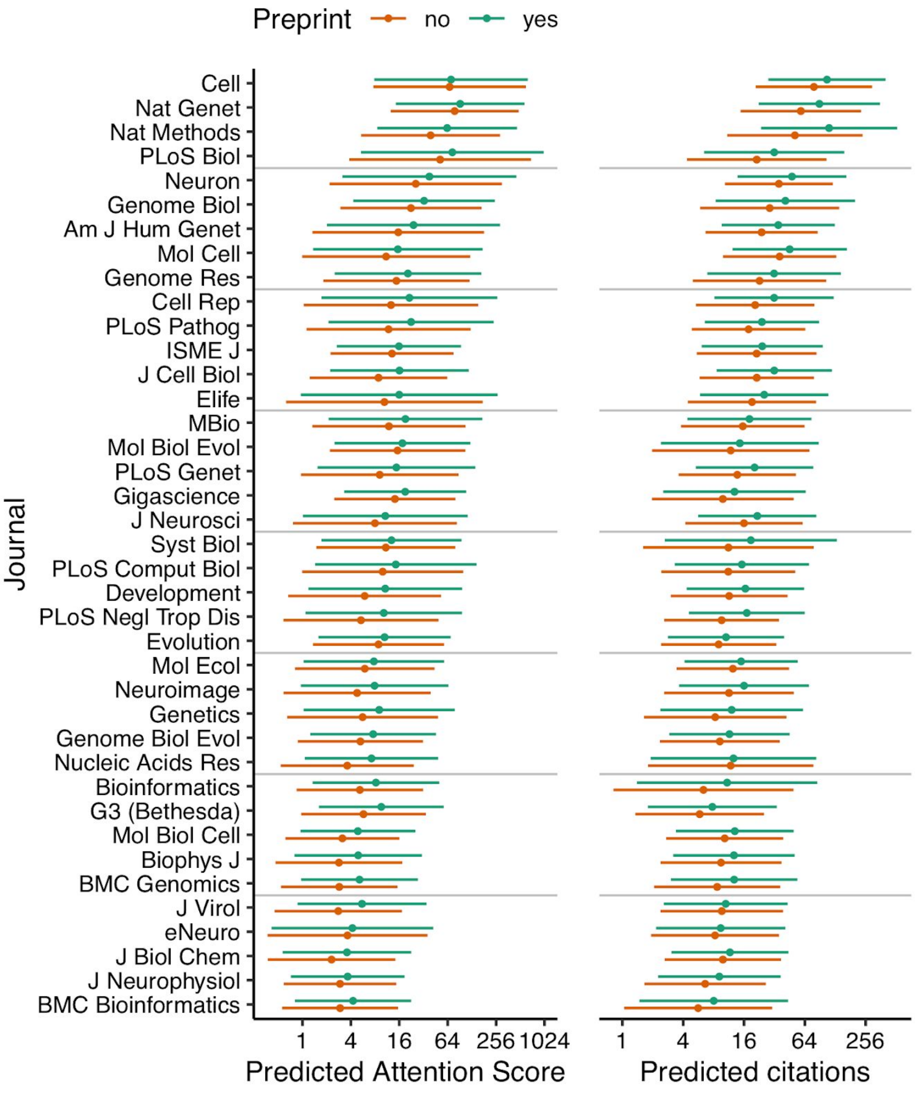


Absolute effect size of having a preprint, by metric and journal. The plots were generated identically to Fig. 1, except they show 95% prediction intervals instead of 95% confidence intervals. Confidence intervals represent uncertainty in the population mean, whereas prediction intervals represent uncertainty in an individual observation. Thus, prediction intervals show the article-to-article variation in Attention Score and citations, even when all variables in the model are fixed.

Figure S9

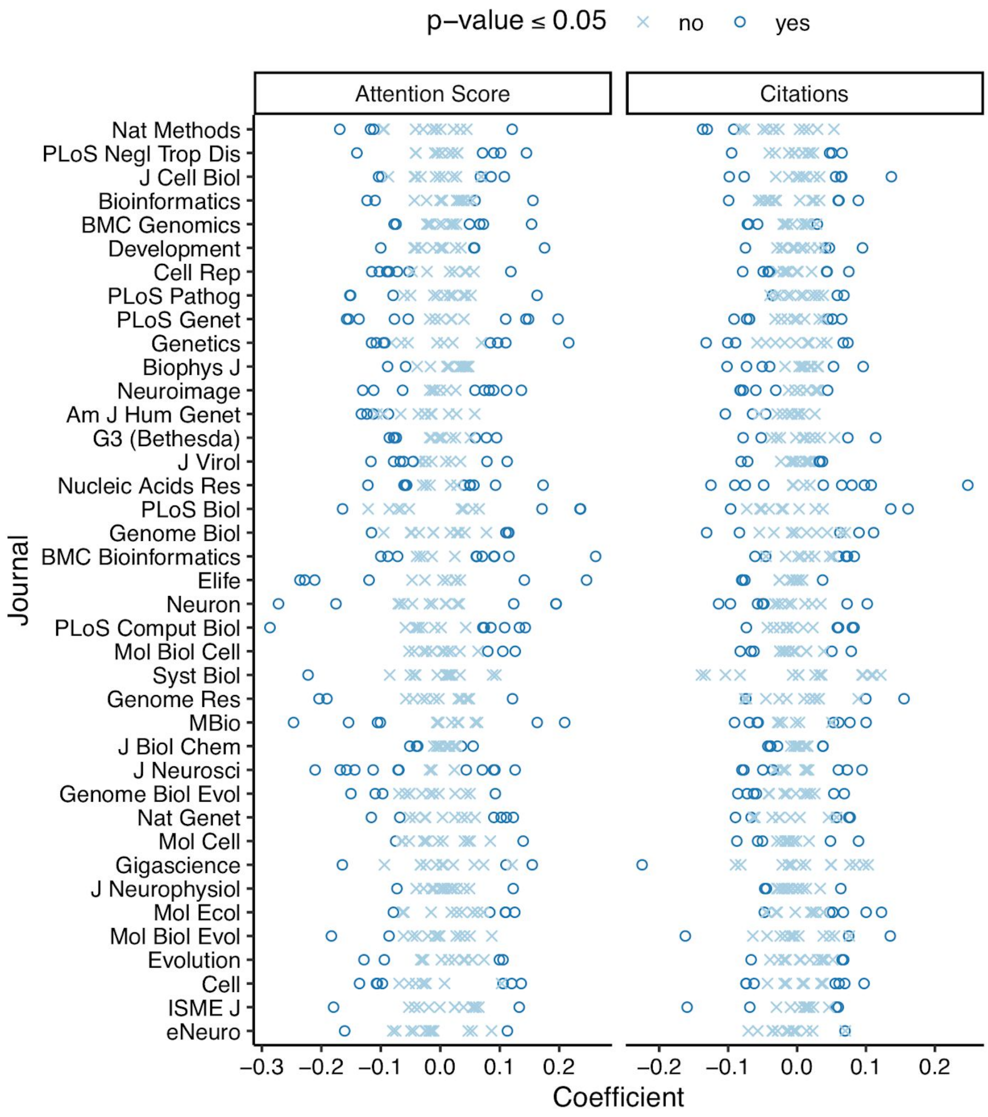


Associations of MeSH term PCs with Attention Score and citations in each journal, based on model coefficients from log-linear regression. P-values are not adjusted for multiple testing.

Figure S10

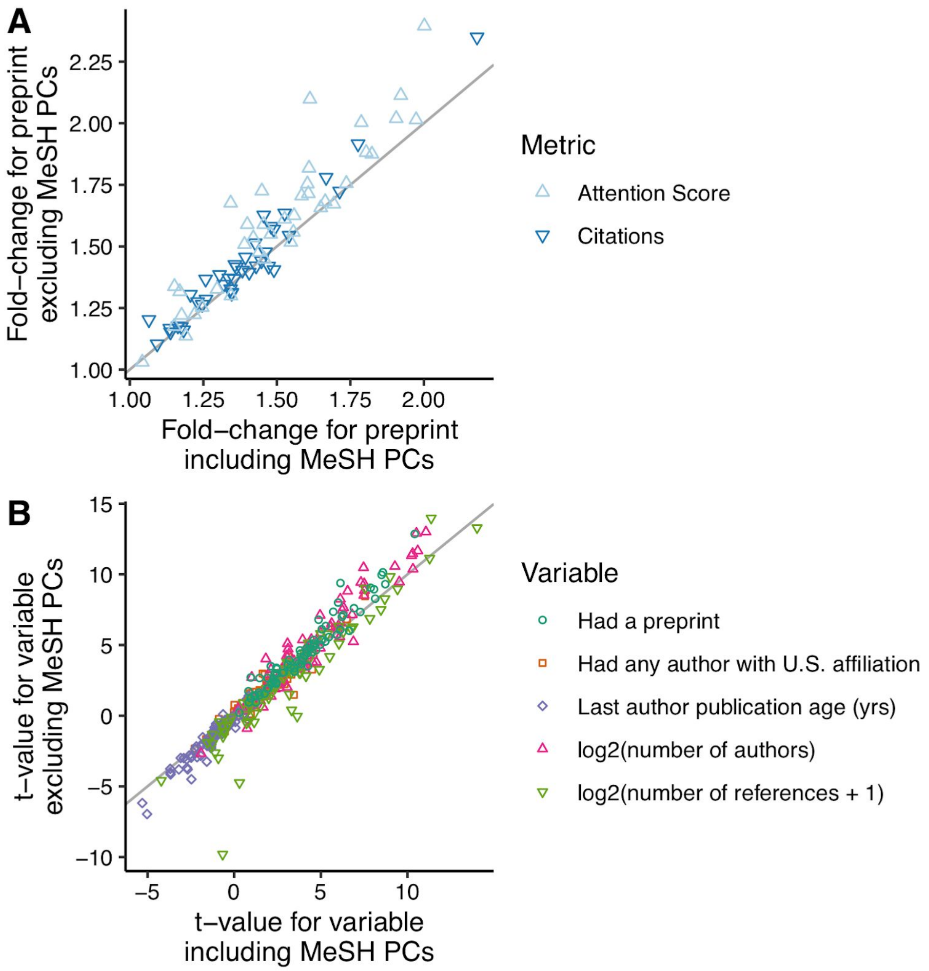


Comparing model fits with and without MeSH term PCs, in terms of **(A)** fold-change (i.e., exponentiated coefficient) for preprint status and **(B)** t-value for each of five variables. Each point represents a journal-metric pair.
